# *Ms4a4a* deficiency ameliorates plaque pathology in a mouse model of amyloid accumulation

**DOI:** 10.1101/2025.03.06.641917

**Authors:** Emma P. Danhash, Anthony C. Verbeck, Daniel Western, Andrea S. Díaz-Pacheco, Grant Galasso, Shih-Feng You, Collin Joseph Nadarajah, Savannah Tiemann Powles, Guangming Huang, Erik S. Musiek, Jasmin Herz, Abhirami K. Iyer, John Cirrito, Carlos Cruchaga, Celeste M. Karch

## Abstract

**Background:** Genome-wide association studies for Alzheimer disease (AD) risk have identified a number of genes enriched in microglia, including *MS4A4A*. Common variants in *MS4A4A* influence AD risk, *MS4A4A* expression, TREM2 signaling, and a specific microglial transcriptional state, though the exact role of MS4A4A in AD remains unclear.

**Methods:** Using a mouse model of amyloid beta (Aβ) accumulation (5xFAD), we examined the impact of *Ms4a4a* loss on Aβ pathology.

**Results:** Before Aβ accumulation, *Ms4a4a* loss reduces steady-state Aβ levels and shortens Aβ half-life in brain interstitial fluid. In aged 5xFAD *Ms4a4a-*deficient mice, plaques are more compact with reduced overall plaque burden. Microglia lacking *Ms4a4a* are more pro-inflammatory and produce more MMP-9, which may promote degradation of Aβ and Aβ fibrils. Human subjects that carry a variant near *MS4A4A* (rs1582763) that confers resilience to AD also exhibit significantly elevated levels of MMP-9 in their cerebrospinal fluid.

**Conclusions:** Together, our results suggest that loss of *Ms4a4a* improves Aβ pathology by altering Aβ clearance, offering insights for therapeutic interventions in AD.

## Background

Genome wide association studies have begun to uncover the complex genetic architecture of Alzheimer’s disease (AD). AD risk loci expression is enriched in microglia [1–9]. Common variants near the *MS4A* gene locus have been associated with AD risk [8, 10] and levels of soluble TREM2 (sTREM2) in the cerebrospinal fluid (CSF) [11, 12]. However, the mechanism by which MS4A4A contributes to AD remains unknown.

MS4A4A is a tetraspan protein enriched in myeloid cells, including microglia [13]. In the periphery, MS4A4A has been proposed to facilitate signaling through recycling of KIT, promote immune response via co-localization with TREM2, and contribute to an anti-inflammatory response [14–18]. In the brain, *MS4A4A* is expressed in a specific microglia transcriptional state that is defined by the expression of anti-inflammatory cytokines in human brains [19, 20]. AD risk genes, including *TREM2*, *CD33*, and *PLCG2* have been shown to alter amyloid accumulation by modifying microglia recruitment and phagocytosis of Aβ [21–25]. Yet, microglia play many critical roles in the brain that may mediate AD pathophysiology. It is unknown whether AD risk genes consistently impact a subset of microglia functions or whether different AD risk genes act on distinct microglia processes.

Here, we sought to define the role of *Ms4a4a* in a model of amyloid pathology (5xFAD). We found that loss of *Ms4a4a* decreases basal Aβ concentration and increases Aβ turnover in the brain interstitial fluid (ISF) prior to plaque accumulation. *Ms4a4a* loss results in a reduced plaque burden in aged mice, with plaques in *Ms4a4a*-deficient mice being more compact.

Interestingly, these phenotypes are not driven by altered microglial recruitment to plaques, as has been described for other AD risk genes like *TREM2*, but instead they are driven by a shift to a pro-inflammatory state and the production of more Aβ-degrading enzyme, MMP-9. In human subjects, an AD resilience variant near the *MS4A4A* gene locus (rs1582763) is associated with elevated MMP-9 levels in the cerebrospinal fluid (CSF). Together, this study suggests that *Ms4a4a* deficiency improves Aβ pathology by altering Aβ degradation via MMP-9.

## Results

### Ms4a4a deficiency in a mouse model of amyloid accumulation

To begin to understand how the AD risk gene, *Ms4a4a,* contributes to AD pathophysiology, guide RNAs (gRNAs) were designed to target intronic regions 5’ to exon 1 and 3’ relative to the stop codon in exon 7 of *Ms4a4a* (**Figure 1A**). Mice with one copy of *Ms4a4a* deleted globally *(Ms4a4a* ^+/-^) were crossed with 5xFAD mice [26], resulting in the generation of our experimental cohort of 5xFAD *Ms4a4a*^+/+^ (5xFAD 4A-WT) and 5xFAD *Ms4a4a*^-/-^ (5xFAD 4A-KO) mice (**Figure 1B**). Quantitative PCR (qPCR) of brain tissue demonstrated that *Ms4a4a* mRNA was absent from 6-month-old 5xFAD 4A-KO mice (**Figure 1C**). *Ms4a4a* is a member of the *Ms4a* super family on chromosome 19 that includes *Ms4a4b*, *Ms4a4c*, *Ms4a4d*, *Ms4a2*, *Ms4a6b*, *Ms4a6c*, *Ms4a6d*, and *Ms4a7* [27, 28]. To determine whether *Ms4a4a* deletion impacts expression of other *Ms4a* family members, transcript levels of *Ms4a4b*, *Ms4a4c*, *Ms4a4d*, *Ms4a2*, *Ms4a6b*, *Ms4a6c*, *Ms4a6d*, and *Ms4a7* were measured by qPCR in brain tissue from 5xFAD 4A-WT and 5xFAD 4A-KO mice (**Figure 1D**). *Ms4a* family genes were expressed at similar levels in 5xFAD 4A-WT and 5xFAD 4A-KO mice (**Figure 1D**). Thus, *Ms4a4a* deletion does not impact the expression of other *Ms4a* family members, and subsequent phenotypes are driven by the loss of *Ms4a4a* rather than broad dysregulation of the *Ms4a* locus.

**Figure 1.**
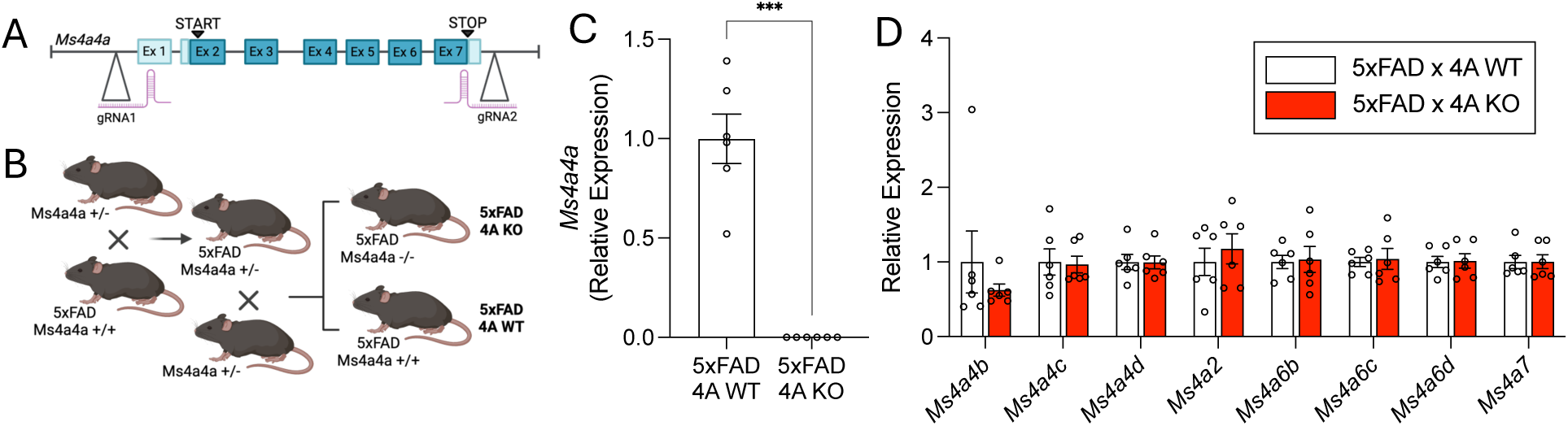
Generation of 5xFAD *Ms4a4a* deficient mice. A. Gene targeting strategy for CRISPR/Cas9-mediated knockout of *Ms4a4a*. B. Breeding scheme for the generation of an experimental cohort of 5xFAD *Ms4a4a*-WT and 5xFAD *Ms4a4a*-KO mice. C. Quantification of *Ms4a4a* levels in 5xFAD 4A-WT and 5xFAD 4A-KO mice reveals the absence of *Ms4a4a* expression in 5xFAD 4A-KO mice (n = 6 mice per genotype) Welch’s t-test, ***, p = 0.0005. D. Quantification of *Ms4a* family gene expression reveals similar expression of *Ms4a* family genes between 5xFAD 4A-WT and 5xFAD 4A-KO mice (n = 6 mice per genotype; 2-way ANOVA).

### Ms4a4a loss promotes soluble ***Aβ*** turnover in 5xFAD mice

Aβ, the primary component of amyloid plaques, is generated when amyloid precursor protein (APP) undergoes a series of enzymatic cleavage events [29, 30]. Aβ is primarily generated by neurons and released into the brain interstitial fluid (ISF), which can be monitored in mice using *in vivo* microdialysis [31, 32]. To determine whether loss of *Ms4a4a* alters soluble Aβ prior to plaque accumulation [33], *in vivo* microdialysis was performed in 8-week-old 5xFAD 4A-KO and 5xFAD 4A-WT mice (**Figure 2A**). Steady-state Aβ levels were significantly reduced in 5xFAD 4A-KO ISF compared with 5xFAD 4A-WT ISF controls (**Figure 2B**; p = 4×10^-4^). To determine whether *Ms4a4a* loss alters the ISF Aβ elimination rate (half-life), 5xFAD 4A-KO and 5xFAD 4A-WT mice were treated with a γ-secretase inhibitor to prevent production of new Aβ. ISF was then sampled hourly to calculate the Aβ elimination rate. ISF Aβ elimination was significantly faster in 5xFAD 4A-KO (0.87 hours) compared with 5xFAD 4A-WT (1.16 hours) mice (**Figure 2C**; p = 0.03). Thus, Aβ levels are reduced and Aβ turnover is accelerated in the absence of *Ms4a4a,* prior to pathology.

**Figure 2.**
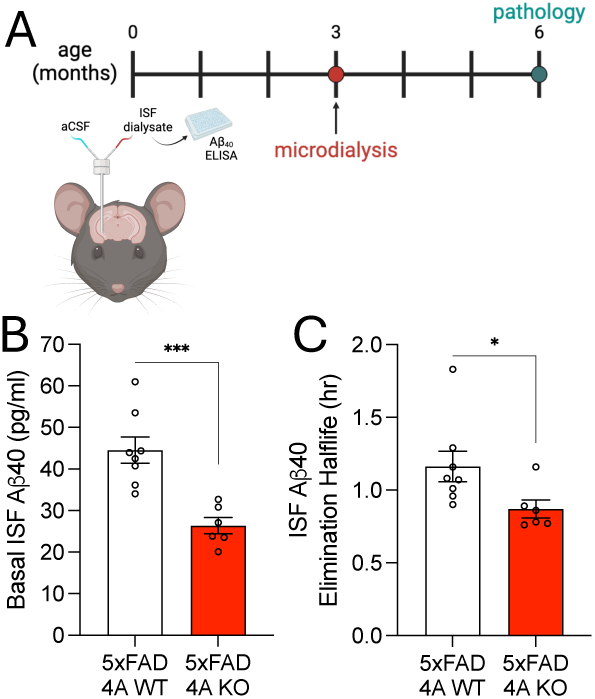
***Ms4a4a* deficiency decreases basal Aβ levels and promotes Aβ turnover in the brain ISF of 5xFAD mice.** A. Schematic depicting experimental timeline and microdialysis set-up. B. Quantification of basal Aβ levels in brain ISF. Welch’s t-test, ***, p=4×10^-4^. C. Quantification of Aβ half-life in brain ISF after treatment with a gamma-secretase inhibitor. Welch’s t-test, *, p=0.03. B-C. 5xFAD 4A-WT n = 9; 5xFAD 4A-KO n = 8; 8-week-old mice.

### Ms4a4a loss reduces plaque burden in 5xFAD mice

The concentration of Aβ in the ISF of young mice has been shown to be highly correlated with the extent of plaque burden in aged mice [34]. Given our findings that young 5xFAD 4A-KO mice produced significantly less ISF Aβ than 5xFAD 4A-WT mice, we hypothesized that 5xFAD 4A-KO mice would display less amyloid pathology with age. To test this hypothesis, 5xFAD 4A-WT and 5xFAD 4A-KO mice were sacrificed at 6 months of age, and brain sections were stained with the HJ3.4 antibody to label total Aβ (**Figure 3A**). Plaque burden, as defined by the area of HJ3.4 coverage, was significantly reduced in the hippocampus and cortex of 5xFAD 4A-KO mice compared with 5xFAD 4A-WT controls (**Figure 3B**; p = 0.01; **Supplemental Figure 1**). The average plaque size was also significantly reduced in the hippocampus of 5xFAD 4A-KO mice (**Figure 3C**; p = 0.05), while the number of plaques per 1000μm^2^ remained similar between the two groups (**Figure 3D**; p = 0.08). Amyloid plaques vary in size, and smaller non-fibrillar plaques are more toxic [35]. To determine the impact of *Ms4a4a* loss on small plaques, plaque frequency was plotted in 349μm^2^ increment bins. 5xFAD 4A-KO mice exhibited significantly fewer smaller plaques (hippocampus: 602-1301μm^2^; cortex: 600-1649μm^2^; **Figure 3E, Supplemental Figure 1**). Together, we show that *Ms4a4a* regulates total plaque burden and clearance of the most toxic forms of plaques.

**Figure 3.**
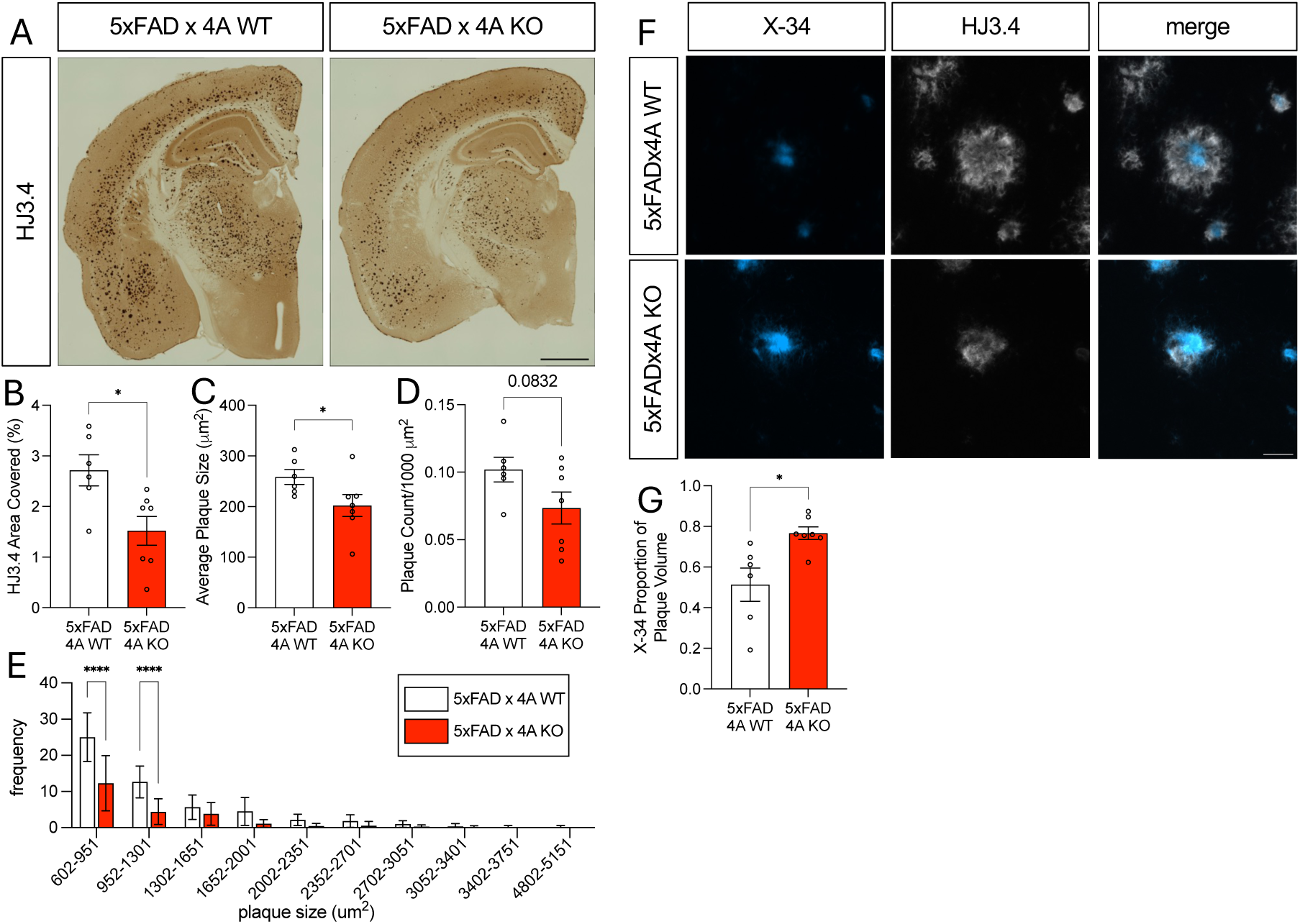
Loss of *Ms4a4a* reduces plaque burden and increases plaque compaction in 5xFAD mice. A. Representative images of immunohistochemistry for Aβ plaques (HJ3.4; scale bar = 1000μm). B. Quantification of HJ3.4 percent area within the hippocampus. Welch’s t-test, *, p = 0.02. C. Quantification of average plaque size within the hippocampus (μm^2^). Welch’s t-test, *, p = 0.05. D. Quantification of average number of plaques/1000μm^2^ within the hippocampus. Welch’s t-test, p = 0.08. E. Quantification of the frequency of plaques binned based on size in μm^2^ within the hippocampus. 2-way ANOVA, ****, p = ˂0.0001. B-E. Each dot represents the mean of 2 brain sections from one animal. F. Representative immunofluorescence confocal images of fibrillar Aβ plaque core (X-34; blue) and total Aβ (HJ3.4; white) in the hippocampus. Scale bar = 20μm. G. Quantification of X-34 proportion relative to total plaque volume. Welch’s t-test; *, p = 0.02. Each dot represents the mean of 2 brain sections from one animal. Data is representative of 5xFAD 4A-WT: n = 6; 5xFAD 4A-KO: n = 7; 6-month-old female mice.

In addition to size, Aβ plaque composition plays a role in plaque toxicity [36]. Aβ plaques are composed of a fibrillar core surrounded by monomeric and oligomeric Aβ [37]. A plaque with a dense fibrillar core (i.e., more compact) has a lower affinity for soluble Aβ, limiting the formation of neurotoxic regions surrounding plaques [36]. We examined whether there was a change in β-sheet rich, dense fibrillar plaques using the X-34 dye. Fibrillar plaque burden, size, and count were similar in 5xFAD 4A-WT and 5xFAD 4A-KO mice within the hippocampus and cortex (**Supplemental Figure 2**). To assess the impact of *Ms4a4a* loss on plaque composition, brain sections were co-stained with X-34 (β-sheet-rich, dense core) and HJ3.4 (total Aβ; **Figure 3F**). Calculating the proportion of X-34 volume relative to total plaque volume revealed that plaques were significantly more compact in the hippocampus of 5xFAD 4A-KO mice than their 5xFAD 4A-WT counterparts (p = 0.02; **Figure 3G**). In the cortex, we observed a similar trend towards more compact plaques in 5xFAD 4A-KO mice; however, the effect was not statistically significant (p = 0.12; **Supplemental Figure 3**). Thus, *Ms4a4a* regulates plaque composition in the hippocampus of 5xFAD mice.

### Ms4a4a loss does not alter microglial recruitment in 5xFAD mice

*Ms4a4a* is enriched in myeloid lineage cells, including microglia [8, 19, 38]. In the central nervous system (CNS), microglia are responsible for phagocytosing and degrading Aβ, clearing it from the parenchyma, and limiting amyloid accumulation [39–44]. Modulation of microglia-enriched AD risk genes such as *Trem2*, *CD33, Inpp5d,* and *Plcg2* have been shown to regulate microglial engulfment of Aβ plaques [21–25]. Thus, we sought to determine whether *Ms4a4a* loss results in altered microglial recruitment and reactivity around plaques in 5xFAD mice. Brain sections from 6-month-old 5xFAD 4A-WT and 5xFAD 4A-KO mice were stained for Aβ plaques (X-34), total microglia (Iba1), and activated microglia (CD68; **Figure 4A**, hippocampus; **Supplemental Figure 4A**, cortex). Iba1-positive microglial recruitment within 15μm of a X-34-positive plaque was similar between 5xFAD 4A-WT and 5xFAD 4A-KO mice (p = 0.65; **Figure 4B**, hippocampus; **Supplemental Figure 4B**, cortex). CD68-positive microglia, representing activated microglia, within 15μm of a X-34-positive plaque were also similar between 5xFAD 4A-WT and 5xFAD 4A-KO mice (p = 0.64; **Figure 4C**, hippocampus; **Supplemental Figure 4C**, cortex). Thus, *Ms4a4a* does not regulate microglial recruitment to plaques.

**Figure 4.**
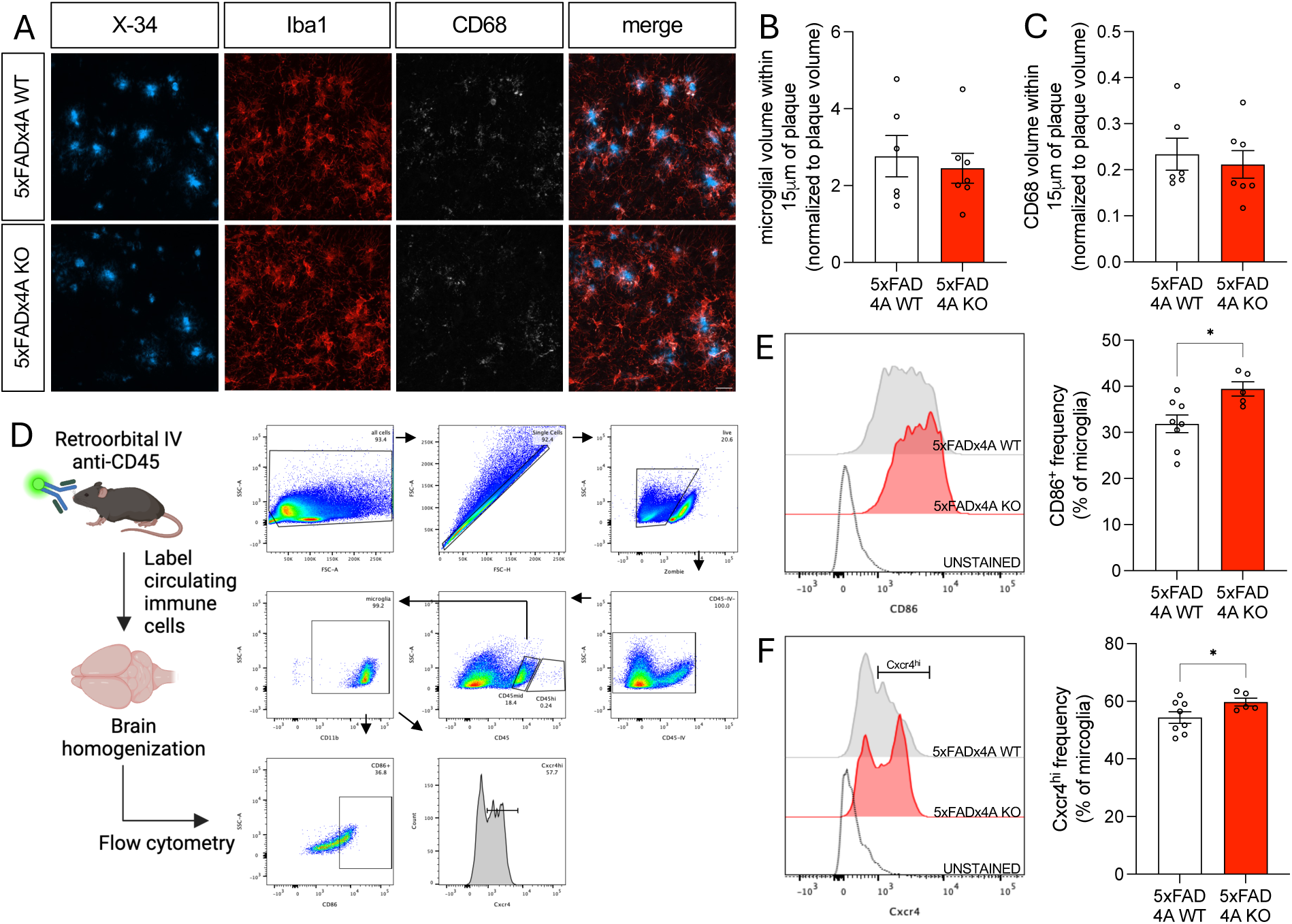
Evaluating the impact of *Ms4a4a* loss on microgliosis in 5xFAD mice. A. Representative confocal images of Aβ plaques (X-34; blue), total microglia (Iba1; red), and a reactive microglia marker (CD68; white; scale bar = 20μm) in the hippocampus. B. Quantification of Iba1-positive microglial volume within 15μm of plaques in 5xFAD 4A-WT and 5xFAD 4A-KO mice in the hippocampus. Welch’s t-test; p = 0.30. C. Quantification of CD68 volume within 15μm of plaques in 5xFAD 4A-WT and 5xFAD 4A-KO mice. Welch’s t-test; p = 0.12. Data is representative of 5xFAD 4A-WT: n = 6; 5xFAD 4A-KO: n = 7. D. Diagram. Mice were retro-orbitally injected with an anti-CD45 antibody to label circulating immune cells, then brains were extracted and homogenized. Flow cytometry was performed. Microglia were gated then analyzed for CD86 and Cxcr4 expression. E. Representative histogram and quantification of CD86 frequency of expression in microglia from 5xFAD 4A-WT (gray) and 5xFAD 4A-KO (red) mice. Welch’s t-test, *, p = 0.0108. F. Representative histogram and quantification of Cxcr4^hi^ frequency of expression in microglia from 5xFAD 4A-WT (gray) and 5xFAD 4A-KO (red) mice. Welch’s t-test, *, p = 0.049. Data is representative of 5xFAD 4A-WT: n = 8; 5xFAD 4A-KO: n = 5; 6-month-old female mice.

To determine whether *Ms4a4a* loss impacts overall microgliosis or astrogliosis, we measured Iba1 and Gfap across the brain. Microgliosis, measured as the percent area of Iba1 staining in the hippocampus and cortex, was similar in 5xFAD 4A-WT and 5xFAD 4A-KO mice (**Supplemental Figure 5A-E**). Total Iba1 intensity, a proxy for microglial activation, was also similar in the hippocampus and cortex of 5xFAD 4A-WT and 5xFAD 4A-KO mice (**Supplemental Figure 5A-E**). Similarly, the degree of astrogliosis, measured as the percent area and intensity of GFAP in the hippocampus or cortex, was similar in 5xFAD 4A-WT and 5xFAD 4A-KO mice (**Supplemental Figure 5F-J**). Thus, *Ms4a4a* does not alter gliosis.

To determine whether *Ms4a4a* loss alters the inflammatory state of microglia, we measured CD86 and Cxcr4 expression in microglia from the whole brain of 6-month-old 5xFAD 4A-WT and 5xFAD 4A-KO mice (**Figure 4D**). FACS analyses revealed that the frequency of microglia expressing CD86, a pro-inflammatory marker, was significantly increased with loss of *Ms4a4a* (**Figure 4E**; p = 0.01). We also found that in the absence of *Ms4a4a*, there were more microglia expressing high levels of Cxcr4, a chemokine receptor involved in inflammation (**Figure 4F**; p = 0.05). Thus, *Ms4a4a* deficiency impacts the inflammatory state of microglia in 5xFAD mice.

### Loss of Ms4a4a increases MMP-9 secretion

Aβ clearance occurs by several mechanisms. Beyond uptake and degradation by activated astrocytes and microglia, soluble and fibrillar forms of Aβ can be enzymatically degraded by extracellular proteases [45]. A number of proteolytic enzymes have been shown to degrade Aβ, including insulin degrading enzyme (IDE)[46–48], neprilysin (NEP)[49, 50], matrix metalloprotease 9 (MMP-9)[51–53], and Cathepsin B[54, 55]. IDE is enriched in astrocytes and only degrades monomeric Aβ, while NEP is expressed in neurons where it degrades Aβ monomers and small oligomers [47, 51, 54, 56–59]. MMP-9 and Cathepsin B are enriched in microglia and digest monomers, oligomers, and fibrils [48, 51, 54, 57–59].

To determine whether *Ms4a4a* loss impacts Aβ-degrading enzymes in 5xFAD mice, we measured levels of Aβ-degrading enzymes by immunoblotting brain homogenates from 6-month-old mice (**Figure 5**). MMP-9 protein levels were significantly increased in the RAB-soluble fraction (enriched for secreted proteins) of 5xFAD 4A-KO brains compared with 5xFAD 4A-WT mice (p = 2.7×10^-3^; **Figure 5A-B**). IDE, Neprilysin, and Cathepsin B protein levels, however, were similar in 5xFAD 4A-KO and 5xFAD 4A-WT mice (p = 0.87, p = 0.14, p = 0.80 respectively; **Figure 5A-B**). We next sought to determine whether Aβ-degrading enzymes were altered in the RIPA-soluble fraction, which contains membrane-bound proteins. While MMP-9, IDE, and NEP levels remain similar (p = 0.98, p = 0.43, p = 0.87, respectively), Cathepsin B levels were significantly reduced in 5xFAD 4A-KO mice compared to 5xFAD 4A-WT mice (**Figure 5C-D**; p = 0.02, respectively). The mRNA expression of these enzymes was similar in 5xFAD 4A-KO and 5xFAD 4A-WT mice, suggesting that *Ms4a4a* acts on Aβ-degrading enzymes at the protein level (**Figure 5E**). Thus, *Ms4a4a* alters microglia-enriched Aβ-degrading enzymes MMP-9 and Cathepsin B, enzymes that degrades Aβ oligomers and fibrils.

**Figure 5.**
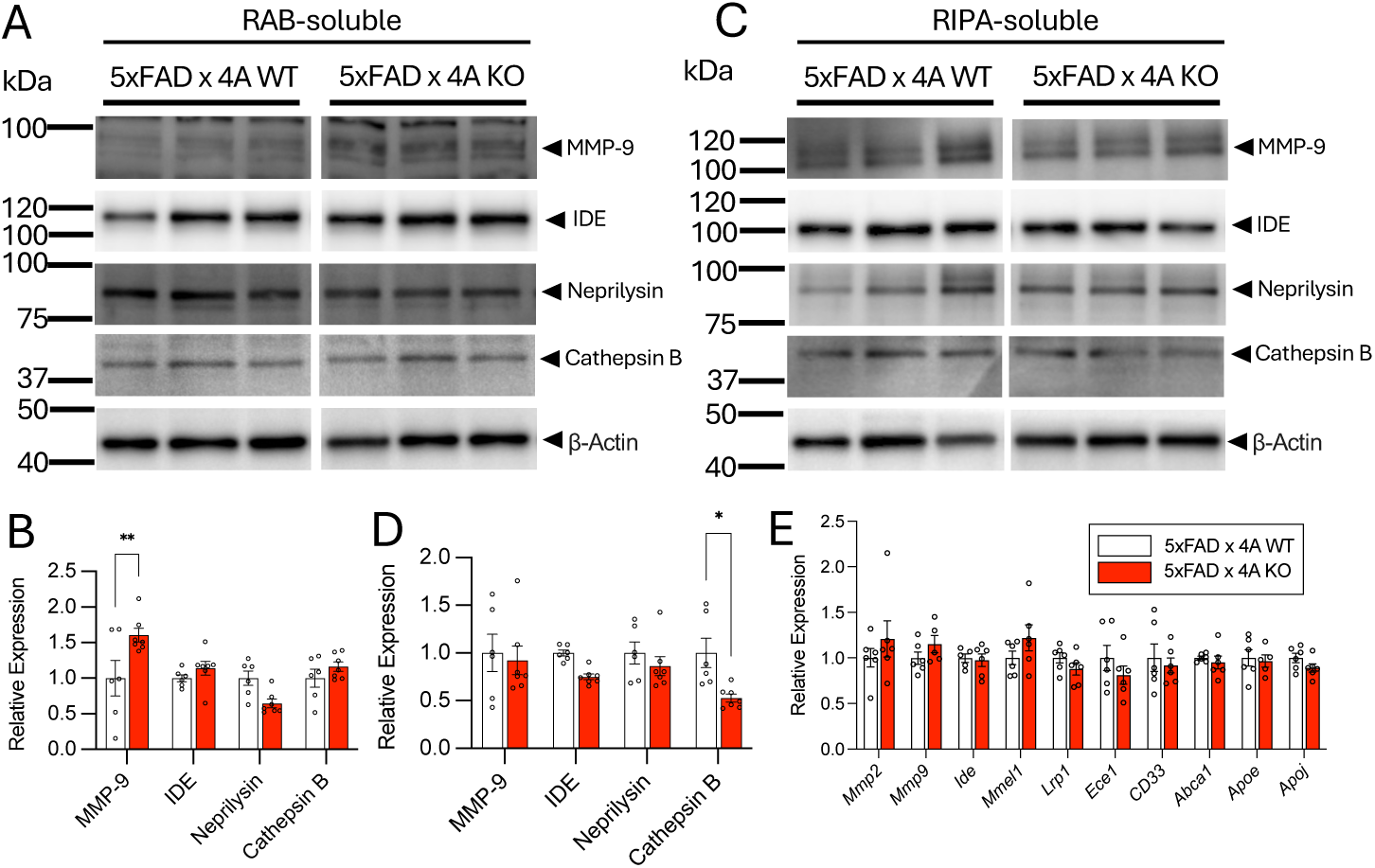
MMP-9 secretion is increased in 5xFAD mice lacking *Ms4a4a*. A. Immunoblot assessing expression of Aβ-degrading enzymes in RAB-soluble protein fraction in 5xFAD 4A-WT and 5xFAD 4A-KO mice. B. Quantification of Aβ-degrading enzyme levels in RAB-soluble protein fraction. 2-way ANOVA; **, p = 2.7×10^-3^). C. Immunoblot of Aβ-degrading enzymes in the RIPA-soluble protein fraction in 5xFAD 4A-WT and 5xFAD 4A-KO mice. D. Quantification of Aβ-degrading enzyme levels in RIPA-soluble protein fraction. 2-way ANOVA; *, p = 0.02). E. Quantification of amyloid-degrading enzyme gene expression reveals similar expression between 5xFAD 4A-WT and 5xFAD 4A-KO mice (2-way ANOVA). Data is representative of 5xFAD 4A-WT: n = 6; 5xFAD 4A-KO: n = 7; 6-month-old female mice.

### AD resilience allele in *MS4A4A* is associated with increased CSF MMP-9 levels

Together, our findings that *Ms4a4a* loss reduces plaque burden and promotes formation of more compact, less toxic plaques, points to a protective phenotype in 5xFAD mice. The minor allele (A; protective) of a common variant near the *MS4A4A* locus (rs1582763) has been associated with reduced risk for AD [11, 60]. To determine whether minor allele carriers of rs1582763 mimic features of the protective phenotypes in the mouse model, we analyzed a cohort of 3,506 subjects with genetic and CSF proteomic data [61]. We found that minor allele (A; protective) carriers had significantly higher levels of CSF MMP-9 than major allele (G) carriers (**Figure 6A**). Interestingly, there is an independent common variant in *MS4A4A* (rs6591561) that confers increased risk for AD [11, 62]. Minor allele carriers (G; risk) of the AD risk variant in *MS4A4A* exhibit significantly lower levels of CSF MMP-9 (**Figure 6B**). Thus, MS4A4A may contribute to AD resilience via modulation of the Aβ-degrading enzyme MMP-9.

**Figure 6.**
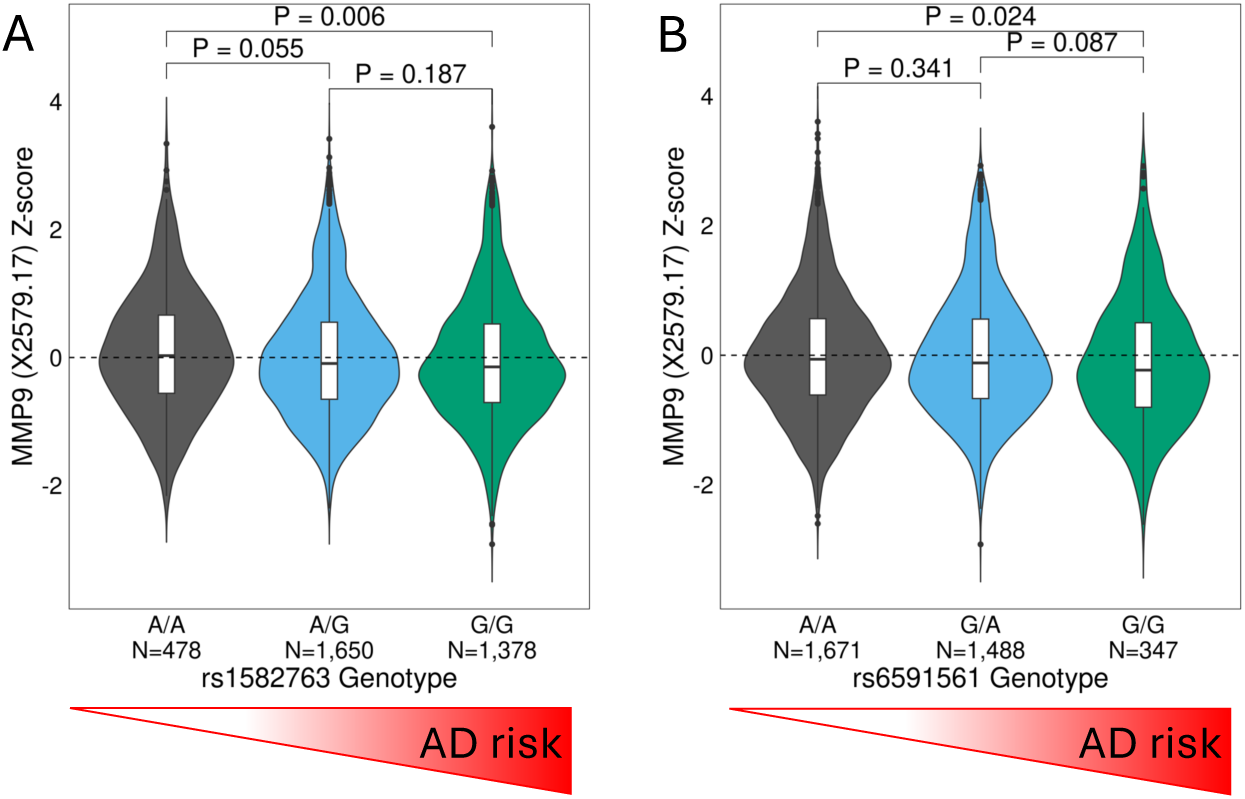
***MS4A4A* variants associated with altered MMP-9 levels in human CSF.**A. Violin plot of MMP-9 levels based on rs1582763 genotype. A, minor allele. Minor allele is associated with reduced AD risk. B. Violin plot of MMP-9 levels based on rs6591561 genotype. G, minor allele. Minor allele is associated with increased AD risk. P-values were obtained using a 2-way Wilcoxon Rank-Sum test.

## Discussion

Common variants in or near *MS4A4A*, a gene implicated in peripheral immune response, are associated with AD risk; however, the role of MS4A4A in AD pathogenesis remains poorly understood [5, 8, 10]. Our study shows that loss of *Ms4a4a* improves pathology in a mouse model of amyloid accumulation. *Ms4a4a* deficiency results in lower basal Aβ levels and faster Aβ turnover in the brain ISF of young mice. This, in turn, contributes to fewer small plaques and increased plaque compaction in aged mice. *Ms4a4a* loss enhances the inflammatory state of microglia without altering recruitment to plaques or overall gliosis. *Ms4a4a* also increases the microglia-enriched, Aβ-fibril-degrading enzymes MMP-9 and decreases Cathepsin B in 5xFAD 4A-KO mice. Consistent with our discovery that *Ms4a4a* loss is protective in the amyloid phase, carriers of a variant that reduces AD risk near *MS4A4A*, rs1582763, have elevated CSF MMP-9 levels, while carriers of an independent variant in *MS4A4A* that increases AD risk, rs6591561, have reduced CSF MMP-9 levels. Together, these results suggest that *MS4A4A* plays a role in mediating amyloid pathobiology by enhancing Aβ clearance.

Plaque accumulation in aged mice corresponds with Aβ concentration in younger mice [34]. We show that *Ms4a4a* is a regulator of ISF Aβ levels and Aβ turnover in a mouse model of plaque pathology. Loss of *Ms4a4a* in 5xFAD mice results in a decrease in ISF Aβ basal levels and half-life. The accumulation of aggregation-prone, clearance-resistant fibrillar Aβ is believed to drive plaque formation and AD and decreased plaque burden is believed to be protective in disease pathogenesis [30, 63]. In line with this, we show that *Ms4a4a-*deficiency results in decreased plaque burden and reduced average plaque size in 6-month-old mice. The reduction in plaque burden is driven by changes in the proportion of small plaques observed in the brains of *Ms4a4a*-deficient 5xFAD mice. This finding is consistent with a number of other microglia-enriched AD risk and resilience genes, including *CD33* and *Inpp5d* [21, 25]. Our findings suggest that loss of *Ms4a4a* is protective against amyloid accumulation. We also find that plaques in the hippocampus of *Ms4a4a-*deficient 5xFAD mice are more “compact” with a dense, fibrillar core surrounded by less non-fibrillar Aβ. Non-fibrillar Aβ is thought to be more toxic, as it has greater potential to seed new plaques and serves as a site of increased neuritic dystrophy [36, 64]. Our finding of more compact Aβ plaques upon *Ms4a4a* loss is consistent with a less toxic environment. Thus, our mouse model of *Ms4a4a* deficiency in the context of plaque pathology appears to capture aspects of resilience in human subjects carrying a common variant near *MS4A4A* (rs1582763) [11, 19, 20].

Slowing of Aβ clearance contributes to late onset AD [65]. Thus, increased clearance of Aβ can result in decreased plaque accumulation thereby decreasing AD pathophysiology.

Microglia, the CNS cell type enriched in *Ms4a4a* expression [13], play an elemental role in phagocytosing and degrading Aβ [39–44]. Loss of *Ms4a4a* fails to alter microglial recruitment in proximity to plaques, as has been described for other AD risk genes like *TREM2* [22]. Instead, we discovered that expression of the pro-inflammatory markers CD86 and Cxcr4 is increased in *Ms4a4a-*deficient microglia in 5xFAD mice. These observations are in line with previous findings which showed that stimulation of peripheral macrophages with IL-4 promotes an MS4A4A-mediated anti-inflammatory, tissue-repair response and that MS4A4A promotes M2 (anti-inflammatory) macrophage polarization in the context of tumor growth [17, 66]. Together, these findings suggest that loss of *Ms4a4a* function results in a more pro-inflammatory polarization of microglia which promotes plaque clearance.

Aβ clearance is also regulated by Aβ degrading enzymes. We found elevated levels of MMP-9, a microglia-enriched [59], zinc-dependent metalloprotease, in *Ms4a4a*-deficient 5xFAD mice. MMP-9 has been shown to degrade not only soluble, but also fibrillar Aβ *in vitro* and *in situ*, with increased activity around compact plaques, generating non-toxic, non-fibrillogenic protein fragments [51–54]. Additionally, overexpression of MMP-9 in 5xFAD mice was found to decrease oligomeric Aβ levels [67]. We propose that the increase of MMP-9 in 5xFAD *Ms4a4a*-KO mice contributes to the reduced plaque pathology observed, consistent with the role of MMP-9 in the clearance of oligomeric and fibrillar Aβ [5, 30]. Furthermore, we discovered that human carriers of a protective variant near *MS4A4A* (rs1582763; A) have significantly elevated levels of CSF MMP-9. rs1582763 is associated with reduced AD risk and delayed age-at-onset, thus offering protection against AD [5, 10, 11, 68]. Carriers of an *MS4A4A* variant associated with increased AD risk (rs6591561) [5, 10, 11, 68], exhibited the opposite effect, whereby CSF MMP-9 levels were significantly reduced in minor allele carriers (G). Our observation of opposing effects of *MS4A4A* variants on pathophysiologic features of disease makes MS4A4A a promising therapeutic target. Together, we demonstrate that the protective effects of Ms4a4a loss in the 5xFAD mouse model are replicated in individuals with an AD protective allele.

Studies in models of amyloid accumulation deficient in microglia-enriched AD risk genes have shown a robust impact on microglial function and reactivity. Microglia are the primary cell type in the brain that express *Ms4a4a* [13]. Loss of *Trem2* in mouse models of amyloid accumulation (5xFAD and APP/PS1) results in exacerbated plaque pathology, more diffuse plaques, increased neuritic dystrophy, and reduced microglial recruitment towards plaques [22, 23, 69, 70]. CD33 inhibits phagocytosis of Aβ42 *in vitro*, and *CD33* deficiency in a mouse model of amyloid accumulation results in reduced plaque burden [21]. In *Ms4a4a*-deficient 5xFAD mice, we identify another mechanism by which microglia function is altered in a manner that modifies amyloid pathology: a global shift towards a more pro-inflammatory microglial activation state. Placing our findings in the context of the growing literature of functional characterization of AD risk genes suggests there are multiple pathways by which AD risk genes alter microglia behavior in ways that contribute to disease, thus there are multiple pathways to target therapeutically.

## Conclusions

We discovered that loss of *Ms4a4a* in the 5xFAD background results in decreased plaque pathology and reflects features of resilience against AD pathogenesis. This work highlights the therapeutic potential of targeting *MS4A4A*, and perhaps modulating the expression of MMP-9, to promote plaque clearance and mitigate not only plaque formation, but also the development of subsequent AD pathologies.

## Methods

### Animals

Animal care and surgical procedures were approved by the Animal Studies Committee of Washington University School of Medicine in accordance with guidelines from the United States National Institutes of Health.

Constitutive *Ms4a4a*^-/-^ mice were generated by Alector using CRISPR/Cas9 technology.

Guide RNAs (gRNAs) were designed to target a noncoding region preceding exon 1 (ACCAGATCCAGTCCTTGTAG) and a noncoding region succeeding exon 7 (GGAATAATCTGCCAACTTCC-TGG; **Figure 1A**) and injected into zygotes to generate *Ms4a4a*^+/-^ mice.

To understand the impact of *Ms4a4a* loss on amyloid pathology, 5xFAD (B6.Cg-Tg(APPSwFlLon, PSEN1*M146L*L286V) 6799Vas/Mmjax, Stock number 034848-JAX, MMRRC, The Jackson Laboratory) hemizygous mice were crossed with the *Ms4a4a*^+/-^ mice to generate 5xFAD x *Ms4a4a*^+/-^ mice. Next, 5xFAD x *Ms4a4a*^+/-^ were crossed with *Ms4a4a*^+/-^ mice to obtain 5xFAD x *Ms4a4a*^-/-^ (5xFAD 4A-KO) mice and 5xFAD x *Ms4a4a*^+/+^ (5xFAD 4A-WT) littermates (**Figure 1B**). Genotype was confirmed using primers to determine the presence of *Ms4a4a* and the 5xFAD transgene (**Supplemental Table 1**). Loss of *Ms4a4a* was validated using quantitative PCR (qPCR) (**Figure 1C; Supplemental Table 2**).

### In Vivo Microdialysis

Prior to plaque accumulation (8 weeks of age), hippocampal ISF Aβ levels were quantified in 5xFAD 4A-KO and 5xFAD 4A-WT littermate controls using microdialysis as previously described [31, 71]. A 38-kDa MWCO semi-permeable membrane was used to allow for molecules smaller than this cut-off to diffuse into the probe. The probe was flushed with perfusion buffer at a constant rate (1.0uL/minute), and samples were collected into a refrigerated fraction collector. ISF samples were then assayed by sandwich ELISA (Aβx-40). A mouse monoclonal anti-Aβ40 capture antibody (mHJ2) made in-house was used in conjunction with a biotinylated central domain detection antibody (mHJ5.1) and streptavidin-poly-HRP-40 (Fitzgerald Industries, Acton, 276 MA) [72]. Mice were housed in cages that permit free movement and ad libitum food and water during microdialysis. ISF Aβ was sampled every 60 minutes between hours 5-12 (after microdialysis probe insertion), and concentrations were averaged to determine the baseline ISF Aβ levels in each mouse. At hour 12, mice were administered the γ-secretase inhibitor (20mg/kg) intraperitoneally, which enabled determination of the elimination half-life of ISF Aβ. Aβ half-life was calculated using first-order kinetics. Statistical difference was measured in GraphPad Prism 10 using an unpaired Student’s t-test with Welch’s correction.

### Brain Tissue Preparation

At 6 months of age, female 5xFAD 4A-KO and 5xFAD 4A-WT mice were anesthetized with sodium pentobarbital and perfused with ice cold PBS. Brains were dissected and cut into 2 hemispheres. The right hemisphere was further dissected to isolate the cortex and hippocampus, which were snap frozen on dry ice and stored at-80°C for biochemical analyses.

The left hemisphere was fixed in 4% paraformaldehyde (w/v) for 48 hours followed by 30% sucrose in PBS at 4°C. Coronal sections (40um) were cut on a freezing-sliding microtome, and slices were collected and stored in cryoprotectant solution (0.2 M phosphate-buffered saline, 30% sucrose, and 30% ethylene glycol) at-20°C.

### 3,3’-diaminobenzidine (DAB) Staining for Amyloid Plaque Quantification

Brain sections from each animal were incubated in 0.3% H_2_O_2_ in Tris-buffered saline (TBS) for 10 minutes then blocked in 3% milk in TBS containing 0.25% Triton-X100 (TBS-X) for 30 minutes. To evaluate plaque pathology, tissue was incubated in HJ3.4b antibody (anti-Aβ-1-13; 1.2μg/mL; a generous gift from the Holtzman lab) diluted in 3% milk in TBS-X overnight at 4°C. Sections were washed with TBS-X, then incubated in Vectastain ABC Elite solution (Vector Laboratories, PK-6100) in TBS for 1 hour at room temperature. Sections were washed with TBS then incubated in DAB solution. Sections were then mounted onto slides and cover slipped with Cytoseal XYL (Electron Microscopy Sciences 18009). Brightfield imaging was performed with a Keyence BZ-X810 microscope.

Plaque burden, plaque count, individual plaque size, and average plaque size were analyzed using NIH ImageJ software. The hippocampus and cortex were analyzed separately for each mouse. Plaque burden was expressed as a percentage of the total area for each brain region and averaged across two slices per animal. Plaque count was expressed as the number of plaques per mm^2^ averaged across two slices for each animal. Plaque size was expressed in µm^2^ averaged across two slices for each animal. Analysis of plaque distribution was performed by stratifying total plaque coverage based on size in μm^2^ and the frequency of occurrence in 349um^2^ increments as previously described [73]. Statistical analyses were performed using GraphPad Prism 10. Differences in plaque burden, size, and count were measured using an unpaired Student’s t-test with Welch’s correction. Statistical difference in plaque distribution was measured using a one-way ANOVA.

### Immunofluorescence and X-34 Plaque Staining

Brain sections from each animal were briefly washed with PBS then permeabilized with Phosphate-buffered saline containing 0.25% Triton-X100 (PBS-X) for 30 minutes at room temperature. Sections were then stained with X-34 (10mM stock solution in DMSO) diluted 1:3000 in staining buffer containing 60% PBS and 40% ethanol, pH 10 for 20 minutes at room temperature. Sections were rinsed in a wash buffer containing 60% PBS and 40% ethanol.

Following X-34 plaque staining, sections were blocked and permeabilized in PBS-X containing 10% goat serum for 1 hour at room temperature, then incubated in one of the following antibodies diluted in PBS-X containing 1% goat serum overnight at 4°C: rabbit anti-Iba1 (Wako 019-19741; 1:500), rat anti-CD68 (BioRad MCA1957; 1:2000), chicken anti-GFAP (abcam ab4674; 1:1000), and HJ3.4b anti-Aβ (a generous gift from the Holtzman lab, 1.2ug/mL).

Sections were washed with PBS then incubated in the following secondary antibodies diluted 1:400 in PBS-X for 1 hour at room temperature: goat anti-rabbit AF568 (Invitrogen A11011), goat anti-rat AF647 (Invitrogen A21247), goat anti-chicken AF647 (Invitrogen A32933), streptavidin AF647 (Invitrogen S21374). Sections were washed with PBS, mounted on slides, and cover slipped with Fluoromount-G mounting medium (Invitrogen 00-4958-02).

### Immunofluorescent Microscopy and Quantification

To analyze Iba1 and GFAP coverage and intensity in the hippocampus and cortex, images were acquired using a Keyence BZ-X810 microscope. Laser intensity and exposure times were set for each staining cohort by surveying tissue and selecting appropriate parameters that could remain the same for all samples. While these values varied by antibody, all sections in a staining cohort were imaged under identical conditions at the same magnification.

To determine percent area stained for markers such as Iba1 and GFAP, TIFF image files were opened in ImageJ and converted to 8-bit greyscale files. Images from the same staining cohort, displaying varying fluorescence intensities, were used to determine an optimal threshold value that could capture staining across all samples. The threshold was then consistently applied to all images in the cohort. Gray scale images were generated for the cortex and hippocampus and quantified as percent area stained using the Analyze Particles function.

Two sections per mouse per region were analyzed and averaged.

All confocal images were acquired using a Zeiss LSM980 microscope. To assess plaque composition, z-stack images of individual plaques were imaged at 20x with 2.2x zoom. To evaluate Iba1 and CD68 proximity to plaques, 20x images were taken.

Confocal images were analyzed in Imaris where 3D reconstructions were made. To determine plaque composition, the volumes of HJ3.4 and X-34 were calculated for each plaque. The X-34 proportion of the plaque volume was then calculated in Microsoft Excel (X-34 volume/(X-34 volume + HJ3.4 volume – colocalized volume)). Imaging and quantification were performed in two brain sections per mouse. Within each brain section, 5 images per brain region (e.g. cortex or hippocampus) were analyzed and averaged (10 images per brain region per mouse). Iba1 and CD68 measurements within 15um of a plaque were classified as “proximal to a plaque”. The volumes of all reconstructions determined to be “proximal to a plaque” were added and normalized to the total plaque volume within an image. Two sections per mouse with one image per brain region per section were analyzed and averaged. Statistical difference was measured in GraphPad Prism 10 using an unpaired Student’s t-test with Welch’s correction.

### Serial Protein Extraction

Frozen cortical tissue was weighed and homogenized using a previously described 3-step serial protein extraction protocol (Yamada et al., 2015). Tissue was first homogenized in RAB buffer (20mg tissue per 400uL buffer; GBiosciences 786-91) supplemented with protease inhibitor cocktail (PI; Sigma P8340; 1:500) and phosphatase inhibitor (PPI; PhosSTOP^™^, Roche 04906837001) with the Red Lysis Kit (1.5mL tubes, Next Advance REDE1) using a bullet blender. Samples were centrifuged at 10,000xg for 5 minutes at 4°C to pellet the RAB-insoluble fraction, then the supernatant was ultracentrifuged at 50,000xg for 20 minutes at 4°C using an MLA-130 rotor in a ultracentrifuge (Beckman) to obtain the RAB-soluble protein fraction. The pellet remaining in the beaded tube was then homogenized in RIPA buffer (150mM NaCl, 50mM Tris, 0.5% deoxycholic acid, 1% Triton X-100, 0.1% SDS, 5mM EDTA, 20mM NaF, and 1mM Na_3_VO_4_, pH 8) supplemented with PI and PPI using a bullet blender. Samples were centrifuged at 10,000xg for 5 minutes at 4°C to pellet the RIPA-insoluble fraction, then the supernatant was sonicated at 10% amplitude for 30 seconds and ultracentrifuged at 50,000xg for 30 minutes at 4°C to obtain the RIPA-soluble protein fraction. The pellet remaining in the beaded tube was then homogenized in 70% formic acid (FA) using a bullet blender then centrifuged at 10,000xg for 5 minutes at 4°C. The supernatant was sonicated at 10% amplitude for 30 seconds and ultracentrifuged at 50,000xg for 20 minutes at 4°C to obtain the FA-soluble protein fraction. The FA-soluble fraction was then neutralized with 1M Tris supplemented with PI and PPI by making a 1:20 dilution. All protein fractions were stored at-80°C for biochemical analyses.

### Immunoblotting

To evaluate total protein concentration of mouse brain RAB-and RIPA-soluble protein fractions, a BCA assay (Pierce^™^ BCA Protein Assay Kit, ThermoFisher Scientific 23225) was performed following manufacturer’s recommendations. To perform SDS-PAGE, 5ug of protein was mixed with 4L×LLaemmli sample buffer (Bio-Rad 161-0747) and 10% β-mercaptoethanol and heated at 95°C for 10 minutes. A 4-12% bis-tris gel (NuPAGE) was used. Proteins were transferred to a PVDF membrane. Membranes were blocked in 5% milk in phosphate buffered saline supplemented with 0.1% Tween 20 (PBS-T) for 1 hour at room temperature then probed with the following antibodies diluted in 5% milk in PBS-T overnight at 4°C: rabbit anti-MMP-9 (abcam ab38898; 1:1000), rabbit anti-neprilysin (abcam ab5458; 1:1000), rabbit anti-insulin degrading enzyme (abcam ab133561; 1:1000), rabbit anti-Cathepsin B (Cell Signaling Technology 3383; 1:1000), and rabbit anti-β-Actin (Cell Signaling Technology 4970; 1:8000). Membranes were then washed and incubated with one of the following secondary antibodies diluted in 5% milk in PBS-T for 1 hour at room temperature: mouse anti-rabbit HRP (Jackson ImmunoResearch 211-032-171; 1:5000) and goat anti-rabbit HRP (Cell Signaling Technology 7074; 1:5000). After washing, membranes were developed using Lumigen ECL Ultra (Lumigen TMA-100) on a Bio-Rad Chemidoc Imaging System. Images were analyzed using Bio-Rad Image Lab Software. Statistical differences were measured in GraphPad Prism 10 using an unpaired Student’s t-test with Welch’s correction.

### Quantitative PCR

Frozen cortical tissue was weighed then homogenized in QIAzol Lysis Reagent (20mg tissue/ 700uL QIAzol; Qiagen 79306) using a handheld tissue grinder. QIAzol samples were then subjected to chloroform extraction followed by mixing and centrifugation at 12,000xg for 15 minutes at 4°C. RNA was extracted from the aqueous layer using a RNeasy Mini Kit (Qiagen 74106) according to manufacturer’s instructions. RNA concentration was measured on a Nanodrop spectrophotometer, then cDNA was made using a High-Capacity cDNA Reverse Transcription Kit (Applied Biosystems 4268814). Real-time qPCR was performed with a Taqman gene expression assay or primers (**Supplemental Table 2**) with iTaq Universal SYBR Green Supermix (BioRad 1725121) using a QuantStudio 12k Flex Real-Time PCR System (ThermoFisher Scientific). Measurements were normalized to *Gapdh* for analysis. Statistical differences were measured in GraphPad Prism 10 using a two-way ANOVA.

### CSF marker association with rs1582763 and rs6591591

Cerebrospinal fluid (CSF) measurements of MMP-9 were generated as part of the SOMAscan7k proteomics panel [61]. The dataset consisted of approximately equal numbers of neurologically healthy controls, individuals with Alzheimer’s Disease, and individuals with other forms of dementia (frontotemporal dementia, Lewy body dementia, Parkinson’s disease). Each individual had genomic data generated using either whole-genome sequencing or array-based genotyping. In total, 3,506 CSF samples from eight cohorts were measured using either the SOMAscan7k or SOMAscan5k panel, after filtering based on quality control or genetic relatedness and selection for individuals of European ancestry.

The dataset underwent stringent quality control at both the protein and individual level to ensure high-quality data. After quality control, protein measurements were converted to the log(10) scale and were converted to z-scores with a mean of zero and standard deviation of 1 using the scale() R function.

Genotypes for each individual at rs1582763 and rs6591561 (GRCh38 coordinates chr11:60254475:G:A and chr11:60302703:A:G) were extracted from the full matrix along with z-score normalized measurements of MMP-9 (SOMAscan aptamer ID X2579.17). In order to control for potential confounding variables, MMP-9 protein levels were residualized using the lm() function in R, where z-score levels of the aptamer were treated as the response variable and age, sex, the first ten genetic principal components, and dummy variables combining both cohort and genotyping array (ex. ADNI_OmniEx) were treated as the terms of the model. The residuals were then extracted, and test statistics for the difference in the residuals for each of the three genotypes (A/A, A/G, G/G) were calculated using a two-sided Wilcoxon test as implemented in the wilcox_test() R function in the rstatix package (version 0.7.2). The violin plots were generated using the ggplot2 (version 3.5.1) and ggpubr (version 0.6.0) R packages.

### Tissue dissociation for flow cytometry

At 6 months of age, 5xFAD 4A-KO and 5xFAD 4A-WT mice were anesthetized with isoflurane and retro-orbitally injected with an APC/Cy7-CD45 antibody (100uL; 3mg/mL). 5-minutes after injection, mice were anesthetized with sodium pentobarbital then perfused with ice cold PBS. The brain was harvested, and the cerebellum and olfactory bulbs were removed. Brains were temporarily stored on ice in Dounce Buffer (1X HBSS, 5mM HEPES, 0.5% glucose).

Tissue was processed using a previously described method [74]. All procedures were performed on ice. Using a razor blade, brains were chopped to roughly 1mm^3^ pieces then transferred to a 5mL Dounce homogenizer containing 5mL of Dounce Buffer. Tissue was homogenized with 6-10 full strokes until the solution appeared to be homogenous then passed through a 40μm strainer into 50mL tubes. Samples were centrifuged and resuspended in 30% Percoll with 2mL of 1X DPBS carefully overlaid. Samples were centrifuged at 400xg for 20 minutes with the brake set to 0. The myelin layer and supernatant were removed, and the remaining cell pellet was resuspended in RPMI 1640 and stored on ice until staining for flow cytometry.

### Flow cytometry

Cells were incubated with Zombie Aqua^TM^ dye (Fixable Viability kit, BioLegend 423102; 1:1000) for 20 minutes in 1X PBS then washed with FACS Buffer (1X PBS, 2% FBS). Samples were incubated with Mouse BD Fc Block^TM^ (anti-CD16/CD32; BD Biosciences 553142; 1:10) for 10 minutes then washed with FACS Buffer. Cells were then stained with Pacific Blue-CD45 (BioLegend 103126; 1:200), PE/Cy7-CD11b (BioLegend 101216; 1:200), APC-Trem2 (R&D Systems FAB17291A; 1:50), BV605-CD86 (BD Biosciences 563055; 1:100), PE-P2ry12 (BioLegend 848004; 1:100), FITC-Cxcr4 (BD Biosciences 551967; 1:100) for 20 minutes then washed with FACS buffer. Cells were examined using a BD FACSSymphony A1 Cell Analyzer with 250,000 events collected per sample. Data was analyzed using FlowJo V10.10.0 software.

## List of Abbreviations

## Declarations

## Ethics approval and consent to participate

All participants provided informed consent for their data to be used in this study. The study was approved by the institutional review board of Washington University School of Medicine in St.Louis.

## Consents for publication

The authors declare that they have no competing interests.

## Availability of data and materials

All data generated or analyzed during this study are included in this published article and its supplementary information files.

## Competing interests

The authors have no competing interests to declare.

## Funding

Funding provided by the National Institutes of Health (AG062734 and AG058501), Hope Center for Neurological Disorders, Chan Zuckerberg Initiative (CMK), Thome Memorial Foundation, and UL1TR002345. The recruitment and clinical characterization of Knight ADRC research participants at Washington University were supported by NIH P30AG066444 (JCM), P01AG03991 (JCM), and P01AG026276 (JCM).

## Authors’ contributions

Designed experiments: EPD, CMK. Performed and analyzed experiments: EPD, ACV, DW, ADP, GG, SFY, CJN, STP, GH, ESM, JH, AKI, JC, CC, CMK. Provided funding: CMK. Wrote the manuscript: EPD, CMK. Revised and approved manuscript: EPD, ACV, DW, ADP, GG, SFY, CJN, STP, GH, ESM, JH, AKI, JC, CC, CMK.

## Acknowledgments

We thank Alector for providing the *Ms4a4a* KO mice used in this study. We thank Dalya Rosner and Torri Ball for thoughtful discussions. This work was supported by access to equipment made possible by the Hope Center for Neurological Disorders, the Neurogenomics and Informatics Center, and the Departments of Neurology and Psychiatry at Washington University School of Medicine. Diagrams were generated using BioRender.com.

## Supplemental Figure Legends

**Supplemental Figure 1. Loss of *Ms4a4a* impacts total amyloid pathology in the cortex of 5xFAD mice.** A. Quantification of HJ3.4 percent area of cortex suggests decreased plaque burden in 5xFAD 4A-KO mice. Welch’s t-test, *, p = 0.04. B. Quantification of HJ3.4-positive average plaque size reveals similar sized plaques in both 5xFAD 4A-WT and 5xFAD 4A-KO mice. Welch’s t-test. C. Quantification of HJ3.4-positive average number of plaques/1000μm^2^ suggests similar number of plaques between 5xFAD 4A-WT and 5xFAD 4A-KO mice. Welch’s t-test, *, p = 0.05. D. Quantification of HJ3.4-positive plaque distribution based on size in μm^2^ and the frequency of occurrence reveals decreased number of smaller plaques in 5xFAD 4A-KO mice. 2-way ANOVA, ***, p = 0.0005, ****, p < 0.0001. Data is representative of 5xFAD 4A-WT: n = 6; 5xFAD 4A-KO: n = 7; 6-month-old female mice.

**Supplemental Figure 2. Loss of *Ms4a4a* does not alter fibrillar plaques in 5xFAD mice.** A. Representative image of fibrillar plaques (X-34; blue) in whole tissue sections (scale bar = 1000μm). B-D. Quantification of X-34-positive percent area (B), average plaque size (C), and average plaque count per 1000μm^2^ (D) in the hippocampus. Welch’s t-test, p = 0.13, p = 0.05, p = 0.20, respectively. E-G. Quantification of X-34-positive percent area (E), average plaque size (F), and average plaque count per 1000μm^2^ (G) in the cortex. Welch’s t-test, p = 0.07, p = 0.15, p = 0.07, respectively. Data is representative of 5xFAD 4A-WT: n = 6; 5xFAD 4A-KO: n = 7; 6-month-old female mice. Welch’s t-test.

**Supplemental Figure 3. Loss of *Ms4a4a* does not alter plaque composition in the cortex of 5xFAD mice.** A. Representative confocal images of immunofluorescence for fibrillar Aβ plaque core (X-34; blue) and total Aβ (HJ3.4; white; scale bar = 20μm). B. Quantification of X-34 proportion of total plaque volume in the cortex reveals similar plaque compaction in 5xFAD 4A-WT and 5xFAD 4A-KO mice Welch’s t-test, p = 0.13. Data is representative of 5xFAD 4A-WT: n = 6; 5xFAD 4A-KO: n = 7; 6-month-old female mice.

**Supplemental Figure 4. Microglia recruitment around plaques in the cortex is similar in 5xFAD with and without *Ms4a4a*.** A. Representative confocal images of Aβ plaques (X-34; blue), microglia (Iba1; red), and microglial lysosomes (CD68; white; scale bar = 20um). B. Quantification of microglial volume within 15μm of plaques suggests similar microglial recruitment to plaques in 5xFAD 4A-WT and 5xFAD 4A-KO mice (Welch’s t-test). Welch’s t-test, p = 0.65. C. Quantification of CD68 volume within 15um of plaques reveals similar microglial reactivity to plaques in 5xFAD 4A-WT and 5xFAD 4A-KO mice. Welch’s t-test, p = 0.96. Data is representative of 5xFAD 4A-WT: n = 6; 5xFAD 4A-KO: n = 7; 6-month-old female mice.

**Supplemental Figure 5. Gliosis remains similar in the cortex with loss of*Ms4a4a* in 5xFAD mice.** A. Representative images of microglia (Iba1; red) in whole tissue sections (scale bar = 1000μm). B-C. Quantification of percent area (B) and intensity (C) of Iba1 reveal similar levels of microgliosis in the hippocampus of 5xFAD 4A-WT and 5xFAD 4A-KO mice. Welch’s t-test, p = 0.30, p = 0.23, respectively. D-E. Quantification of percent area (D) and intensity (E) of Iba1 reveal similar levels of microgliosis in the cortex of 5xFAD 4A-WT and 5xFAD 4A-KO mice. Welch’s t-test, p = 0.10, p = 0.07, respectively. F. Representative images of astrocytes (GFAP; white) in whole tissue sections (scale bar = 1000μm). G-H. Quantification of percent area (G) and intensity (H) of GFAP reveal similar levels of astrogliosis in the hippocampus of 5xFAD 4A-WT and 5xFAD 4A-KO mice. Welch’s t-test, p = 0.40, p = 0.20, respectively. I-J. Quantification of percent area (I) and intensity (J) of Iba1 and GFAP reveal similar levels of astrogliosis in the cortex of 5xFAD 4A-WT and 5xFAD 4A-KO mice. Welch’s t-test, p = 0.29, p = 0.14, respectively. Data is representative of 5xFAD 4A-WT: n = 6; 5xFAD 4A-KO: n = 7; 6-month-old female mice.

## Notes

### Competing Interest Statement

The authors have declared no competing interest.

## References

1. Efthymiou AG, Goate AM: Late onset Alzheimer’s disease genetics implicates microglial pathways in disease risk. Mol Neurodegener 2017, 12:43.

2. Guerreiro R, Wojtas A, Bras J, Carrasquillo M, Rogaeva E, Majounie E, Cruchaga C, Sassi C, Kauwe JS, Younkin S, et al: TREM2 variants in Alzheimer’s disease. N Engl J Med 2013, 368:117–127.

3. Jonsson T, Stefansson H, Steinberg S, Jonsdottir I, Jonsson PV, Snaedal J, Bjornsson S, Huttenlocher J, Levey AI, Lah JJ, et al: Variant of TREM2 associated with the risk of Alzheimer’s disease. N Engl J Med 2013, 368:107–116.

4. Karch CM, Goate AM: Alzheimer’s disease risk genes and mechanisms of disease pathogenesis. Biol Psychiatry 2015, 77:43–51.

5. Sims R, van der Lee SJ, Naj AC, Bellenguez C, Badarinarayan N, Jakobsdottir J, Kunkle BW, Boland A, Raybould R, Bis JC, et al: Rare coding variants in PLCG2, ABI3, and TREM2 implicate microglial-mediated innate immunity in Alzheimer’s disease. Nat Genet 2017, 49:1373–1384.

6. Brase L, You S-F, del Aguila J, Dai Y, Novotny BC, Soriano-Tarraga C, Dykstra T, Fernandez MV, Budde JP, Bergmann K, et al: A landscape of the genetic and cellular heterogeneity in Alzheimer disease. medRxiv 2021:2021.2011.2030.21267072.

7. Nott A, Holtman IR, Coufal NG, Schlachetzki JCM, Yu M, Hu R, Han CZ, Pena M, Xiao J, Wu Y, et al: Brain cell type-specific enhancer-promoter interactome maps and disease-risk association. Science 2019, 366:1134–1139.

8. Novikova G, Kapoor M, Tcw J, Abud EM, Efthymiou AG, Chen SX, Cheng H, Fullard JF, Bendl J, Liu Y, et al: Integration of Alzheimer’s disease genetics and myeloid genomics identifies disease risk regulatory elements and genes.Nat Commun 2021, 12:1610.

9. Treusch S, Hamamichi S, Goodman JL, Matlack KE, Chung CY, Baru V, Shulman JM, Parrado A, Bevis BJ, Valastyan JS, et al: Functional links between Abeta toxicity, endocytic trafficking, and Alzheimer’s disease risk factors in yeast.Science 2011, 334:1241–1245.

10. Wightman DP, Jansen IE, Savage JE, Shadrin AA, Bahrami S, Holland D, Rongve A, Borte S, Winsvold BS, Drange OK, et al: A genome-wide association study with 1,126,563 individuals identifies new risk loci for Alzheimer’s disease. Nat Genet 2021, 53:1276–1282.

11. Deming Y, Filipello F, Cignarella F, Cantoni C, Hsu S, Mikesell R, Li Z, Del-Aguila JL, Dube U, Farias FG, et al: The MS4A gene cluster is a key modulator of soluble TREM2 and Alzheimer’s disease risk. Sci Transl Med 2019, 11.

12. Wang L, Nykanen NP, Western D, Gorijala P, Timsina J, Li F, Wang Z, Ali M, Yang C, Liu M, et al: Proteo-genomics of soluble TREM2 in cerebrospinal fluid provides novel insights and identifies novel modulators for Alzheimer’s disease.Mol Neurodegener 2024, 19:1.

13. Mattiola I, Mantovani A, Locati M: The tetraspan MS4A family in homeostasis, immunity, and disease. Trends Immunol 2021, 42:764–781.

14. Liu W, Taso O, Wang R, Bayram S, Graham AC, Garcia-Reitboeck P, Mallach A, Andrews WD, Piers TM, Botia JA, et al: Trem2 promotes anti-inflammatory responses in microglia and is suppressed under pro-inflammatory conditions.Hum Mol Genet 2020, 29:3224–3248.

15. Mattiola I, Tomay F, De Pizzol M, Silva-Gomes R, Savino B, Gulic T, Doni A, Lonardi S, Astrid Boutet M, Nerviani A, et al: The macrophage tetraspan MS4A4A enhances dectin-1-dependent NK cell-mediated resistance to metastasis. Nat Immunol 2019, 20:1012–1022.

16. Zhao X, Sun J, Xiong L, She L, Li L, Tang H, Zeng Y, Chen F, Han X, Ye S, et al: beta-amyloid binds to microglia Dectin-1 to induce inflammatory response in the pathogenesis of Alzheimer’s disease. Int J Biol Sci 2023, 19:3249–3265.

17. Sanyal R, Polyak MJ, Zuccolo J, Puri M, Deng L, Roberts L, Zuba A, Storek J, Luider JM, Sundberg EM, et al: MS4A4A: a novel cell surface marker for M2 macrophages and plasma cells. Immunol Cell Biol 2017, 95:611–619.

18. Cruse G, Beaven MA, Music SC, Bradding P, Gilfillan AM, Metcalfe DD: The CD20 homologue MS4A4 directs trafficking of KIT toward clathrin-independent endocytosis pathways and thus regulates receptor signaling and recycling.Mol Biol Cell 2015, 26:1711–1727.

19. Brase L, You SF, D’Oliveira Albanus R, Del-Aguila JL, Dai Y, Novotny BC, Soriano-Tarraga C, Dykstra T, Fernandez MV, Budde JP, et al: Single-nucleus RNA-sequencing of autosomal dominant Alzheimer disease and risk variant carriers.Nat Commun 2023, 14:2314.

20. You SF, Brase L, Filipello F, Iyer AK, Del-Aguila J, He J, D’Oliveira Albanus R, Budde J, Norton J, Gentsch J, et al: MS4A4A modifies the risk of Alzheimer disease by regulating lipid metabolism and immune response in a unique microglia state. medRxiv 2023.

21. Griciuc A, Serrano-Pozo A, Parrado AR, Lesinski AN, Asselin CN, Mullin K, Hooli B, Choi SH, Hyman BT, Tanzi RE: Alzheimer’s disease risk gene CD33 inhibits microglial uptake of amyloid beta. Neuron 2013, 78:631–643.

22. Ulrich JD, Finn MB, Wang Y, Shen A, Mahan TE, Jiang H, Stewart FR, Piccio L, Colonna M, Holtzman DM: Altered microglial response to Abeta plaques in APPPS1-21 mice heterozygous for TREM2. Mol Neurodegener 2014, 9:20.

23. Yuan P, Condello C, Keene CD, Wang Y, Bird TD, Paul SM, Luo W, Colonna M, Baddeley D, Grutzendler J: TREM2 Haplodeficiency in Mice and Humans Impairs the Microglia Barrier Function Leading to Decreased Amyloid Compaction and Severe Axonal Dystrophy. Neuron 2016, 92:252–264.

24. Tsai AP, Dong C, Lin PB, Oblak AL, Viana Di Prisco G, Wang N, Hajicek N, Carr AJ, Lendy EK, Hahn O, et al: Genetic variants of phospholipase C-gamma2 alter the phenotype and function of microglia and confer differential risk for Alzheimer’s disease.Immunity 2023, 56:2121–2136 e2126.

25. Lin PB, Tsai AP, Soni D, Lee-Gosselin A, Moutinho M, Puntambekar SS, Landreth GE, Lamb BT, Oblak AL: INPP5D deficiency attenuates amyloid pathology in a mouse model of Alzheimer’s disease. Alzheimers Dement 2023, 19:2528–2537.

26. Oakley H, Cole SL, Logan S, Maus E, Shao P, Craft J, Guillozet-Bongaarts A, Ohno M, Disterhoft J, Van Eldik L, et al: Intraneuronal beta-amyloid aggregates, neurodegeneration, and neuron loss in transgenic mice with five familial Alzheimer’s disease mutations: potential factors in amyloid plaque formation.J Neurosci 2006, 26:10129–10140.

27. Ishibashi K, Suzuki M, Sasaki S, Imai M: Identification of a new multigene four-transmembrane family (MS4A) related to CD20, HTm4 and beta subunit of the high-affinity IgE receptor. Gene 2001, 264:87–93.

28. Liang Y, Tedder TF: Identification of a CD20-, FcepsilonRIbeta-, and HTm4-related gene family: sixteen new MS4A family members expressed in human and mouse. Genomics 2001, 72:119-127.

29. Haass C, Kaether C, Thinakaran G, Sisodia S: Trafficking and proteolytic processing of APP. Cold Spring Harb Perspect Med 2012, 2:a006270.

30. Selkoe DJ: Alzheimer’s disease results from the cerebral accumulation and cytotoxicity of amyloid beta-protein. J Alzheimers Dis 2001, 3:75–80.

31. Cirrito JR, May PC, O’Dell MA, Taylor JW, Parsadanian M, Cramer JW, Audia JE, Nissen JS, Bales KR, Paul SM, et al: In vivo assessment of brain interstitial fluid with microdialysis reveals plaque-associated changes in amyloid-beta metabolism and half-life.J Neurosci 2003, 23:8844–8853.

32. Kamenetz F, Tomita T, Hsieh H, Seabrook G, Borchelt D, Iwatsubo T, Sisodia S, Malinow R: APP processing and synaptic function. Neuron 2003, 37:925–937.

33. Jawhar S, Trawicka A, Jenneckens C, Bayer TA, Wirths O: Motor deficits, neuron loss, and reduced anxiety coinciding with axonal degeneration and intraneuronal Abeta aggregation in the 5XFAD mouse model of Alzheimer’s disease.Neurobiol Aging 2012, 33:196 e129–140.

34. Bero AW, Yan P, Roh JH, Cirrito JR, Stewart FR, Raichle ME, Lee JM, Holtzman DM: Neuronal activity regulates the regional vulnerability to amyloid-beta deposition.Nat Neurosci 2011, 14:750–756.

35. Sengupta U, Nilson AN, Kayed R: The Role of Amyloid-beta Oligomers in Toxicity, Propagation, and Immunotherapy. EBioMedicine 2016, 6:42–49.

36. Condello C, Yuan P, Schain A, Grutzendler J: Microglia constitute a barrier that prevents neurotoxic protofibrillar Abeta42 hotspots around plaques.Nat Commun 2015, 6:6176.

37. Walker LC: Abeta Plaques. Free Neuropathol 2020, 1.

38. Silva-Gomes R, Mapelli SN, Boutet MA, Mattiola I, Sironi M, Grizzi F, Colombo F, Supino D, Carnevale S, Pasqualini F, et al: Differential expression and regulation of MS4A family members in myeloid cells in physiological and pathological conditions.J Leukoc Biol 2022, 111:817–836.

39. Bamberger ME, Harris ME, McDonald DR, Husemann J, Landreth GE: A cell surface receptor complex for fibrillar beta-amyloid mediates microglial activation.J Neurosci 2003, 23:2665–2674.

40. Lee CY, Landreth GE: The role of microglia in amyloid clearance from the AD brain.J Neural Transm (Vienna) 2010, 117:949–960.

41. Frackowiak J, Wisniewski HM, Wegiel J, Merz GS, Iqbal K, Wang KC: Ultrastructure of the microglia that phagocytose amyloid and the microglia that produce beta-amyloid fibrils. Acta Neuropathol 1992, 84:225–233.

42. Bolmont T, Haiss F, Eicke D, Radde R, Mathis CA, Klunk WE, Kohsaka S, Jucker M, Calhoun ME: Dynamics of the microglial/amyloid interaction indicate a role in plaque maintenance. J Neurosci 2008, 28:4283–4292.

43. Meyer-Luehmann M, Spires-Jones TL, Prada C, Garcia-Alloza M, de Calignon A, Rozkalne A, Koenigsknecht-Talboo J, Holtzman DM, Bacskai BJ, Hyman BT: Rapid appearance and local toxicity of amyloid-beta plaques in a mouse model of Alzheimer’s disease.Nature 2008, 451:720–724.

44. Hansen DV, Hanson JE, Sheng M: Microglia in Alzheimer’s disease. J Cell Biol 2018, 217:459–472.

45. Yoon SS, Jo SA: Mechanisms of Amyloid-beta Peptide Clearance: Potential Therapeutic Targets for Alzheimer’s Disease. Biomol Ther (Seoul) 2012, 20:245–255.

46. Vekrellis K, Ye Z, Qiu WQ, Walsh D, Hartley D, Chesneau V, Rosner MR, Selkoe DJ: Neurons regulate extracellular levels of amyloid beta-protein via proteolysis by insulin-degrading enzyme. J Neurosci 2000, 20:1657–1665.

47. Qiu WQ, Walsh DM, Ye Z, Vekrellis K, Zhang J, Podlisny MB, Rosner MR, Safavi A, Hersh LB, Selkoe DJ: Insulin-degrading enzyme regulates extracellular levels of amyloid beta-protein by degradation. J Biol Chem 1998, 273:32730–32738.

48. Qiu WQ, Ye Z, Kholodenko D, Seubert P, Selkoe DJ: Degradation of amyloid beta-protein by a metalloprotease secreted by microglia and other neural and non-neural cells.J Biol Chem 1997, 272:6641–6646.

49. Iwata N, Tsubuki S, Takaki Y, Watanabe K, Sekiguchi M, Hosoki E, Kawashima-Morishima M, Lee HJ, Hama E, Sekine-Aizawa Y, Saido TC: Identification of the major Abeta1-42-degrading catabolic pathway in brain parenchyma: suppression leads to biochemical and pathological deposition. Nat Med 2000, 6:143–150.

50. Shirotani K, Tsubuki S, Iwata N, Takaki Y, Harigaya W, Maruyama K, Kiryu-Seo S, Kiyama H, Iwata H, Tomita T, et al: Neprilysin degrades both amyloid beta peptides 1-40 and 1-42 most rapidly and efficiently among thiorphan-and phosphoramidon-sensitive endopeptidases. J Biol Chem 2001, 276:21895–21901.

51. Hernandez-Guillamon M, Mawhirt S, Blais S, Montaner J, Neubert TA, Rostagno A, Ghiso J: Sequential Amyloid-beta Degradation by the Matrix Metalloproteases MMP-2 and MMP-9. J Biol Chem 2015, 290:15078–15091.

52. Yan P, Hu X, Song H, Yin K, Bateman RJ, Cirrito JR, Xiao Q, Hsu FF, Turk JW, Xu J, et al: Matrix metalloproteinase-9 degrades amyloid-beta fibrils in vitro and compact plaques in situ. J Biol Chem 2006, 281:24566–24574.

53. Backstrom JR, Lim GP, Cullen MJ, Tokes ZA: Matrix metalloproteinase-9 (MMP-9) is synthesized in neurons of the human hippocampus and is capable of degrading the amyloid-beta peptide (1-40). J Neurosci 1996, 16:7910–7919.

54. Mueller-Steiner S, Zhou Y, Arai H, Roberson ED, Sun B, Chen J, Wang X, Yu G, Esposito L, Mucke L, Gan L: Antiamyloidogenic and neuroprotective functions of cathepsin B: implications for Alzheimer’s disease. Neuron 2006, 51:703–714.

55. Mackay EA, Ehrhard A, Moniatte M, Guenet C, Tardif C, Tarnus C, Sorokine O, Heintzelmann B, Nay C, Remy JM, et al: A possible role for cathepsins D, E, and B in the processing of beta-amyloid precursor protein in Alzheimer’s disease. Eur J Biochem 1997, 244:414–425.

56. Walsh DM, Klyubin I, Fadeeva JV, Cullen WK, Anwyl R, Wolfe MS, Rowan MJ, Selkoe DJ: Naturally secreted oligomers of amyloid beta protein potently inhibit hippocampal long-term potentiation in vivo. Nature 2002, 416:535–539.

57. Sudoh S, Frosch MP, Wolf BA: Differential effects of proteases involved in intracellular degradation of amyloid beta-protein between detergent-soluble and-insoluble pools in CHO-695 cells. Biochemistry 2002, 41:1091–1099.

58. Howell S, Nalbantoglu J, Crine P: Neutral endopeptidase can hydrolyze beta-amyloid(1-40) but shows no effect on beta-amyloid precursor protein metabolism.Peptides 1995, 16:647–652.

59. Zhang Y, Chen K, Sloan SA, Bennett ML, Scholze AR, O’Keeffe S, Phatnani HP, Guarnieri P, Caneda C, Ruderisch N, et al: An RNA-Sequencing Transcriptome and Splicing Database of Glia, Neurons, and Vascular Cells of the Cerebral Cortex. J Neurosci 2014, 34:11929–11947.

60. Naj AC, Jun G, Beecham GW, Wang LS, Vardarajan BN, Buros J, Gallins PJ, Buxbaum JD, Jarvik GP, Crane PK, et al: Common variants at MS4A4/MS4A6E, CD2AP, CD33 and EPHA1 are associated with late-onset Alzheimer’s disease. Nat Genet 2011, 43:436–441.

61. Western D, Timsina J, Wang L, Wang C, Yang C, Phillips B, Wang Y, Liu M, Ali M, Beric A, et al: Proteogenomic analysis of human cerebrospinal fluid identifies neurologically relevant regulation and implicates causal proteins for Alzheimer’s disease. Nat Genet 2024.

62. Naj AC, Jun G, Reitz C, Kunkle BW, Perry W, Park YS, Beecham GW, Rajbhandary RA, Hamilton-Nelson KL, Wang LS, et al: Effects of multiple genetic loci on age at onset in late-onset Alzheimer disease: a genome-wide association study.JAMA Neurol 2014, 71:1394–1404.

63. Hardy JA, Higgins GA: Alzheimer’s disease: the amyloid cascade hypothesis.Science 1992, 256:184–185.

64. Vander Zanden CM, Wampler L, Bowers I, Watkins EB, Majewski J, Chi EY: Fibrillar and Nonfibrillar Amyloid Beta Structures Drive Two Modes of Membrane-Mediated Toxicity.Langmuir 2019, 35:16024–16036.

65. Mawuenyega KG, Sigurdson W, Ovod V, Munsell L, Kasten T, Morris JC, Yarasheski KE, Bateman RJ: Decreased clearance of CNS beta-amyloid in Alzheimer’s disease.Science 2010, 330:1774.

66. Li Y, Shen Z, Chai Z, Zhan Y, Zhang Y, Liu Z, Liu Y, Li Z, Lin M, Zhang Z, et al: Targeting MS4A4A on tumour-associated macrophages restores CD8+ T-cell-mediated antitumour immunity. Gut 2023, 72:2307–2320.

67. Kaminari A, Giannakas N, Tzinia A, Tsilibary EC: Overexpression of matrix metalloproteinase-9 (MMP-9) rescues insulin-mediated impairment in the 5XFAD model of Alzheimer’s disease. Sci Rep 2017, 7:683.

68. Lambert JC, Ibrahim-Verbaas CA, Harold D, Naj AC, Sims R, Bellenguez C, DeStafano AL, Bis JC, Beecham GW, Grenier-Boley B, et al: Meta-analysis of 74,046 individuals identifies 11 new susceptibility loci for Alzheimer’s disease.Nat Genet 2013, 45:1452–1458.

69. Wang Y, Ulland TK, Ulrich JD, Song W, Tzaferis JA, Hole JT, Yuan P, Mahan TE, Shi Y, Gilfillan S, et al: TREM2-mediated early microglial response limits diffusion and toxicity of amyloid plaques. J Exp Med 2016, 213:667–675.

70. Jay TR, Hirsch AM, Broihier ML, Miller CM, Neilson LE, Ransohoff RM, Lamb BT, Landreth GE: Disease Progression-Dependent Effects of TREM2 Deficiency in a Mouse Model of Alzheimer’s Disease. J Neurosci 2017, 37:637–647.

71. Cirrito JR, Yamada KA, Finn MB, Sloviter RS, Bales KR, May PC, Schoepp DD, Paul SM, Mennerick S, Holtzman DM: Synaptic activity regulates interstitial fluid amyloid-beta levels in vivo. Neuron 2005, 48:913–922.

72. Hettinger JC, Lee H, Bu G, Holtzman DM, Cirrito JR: AMPA-ergic regulation of amyloid-beta levels in an Alzheimer’s disease mouse model. Mol Neurodegener 2018, 13:22.

73. Huynh TV, Liao F, Francis CM, Robinson GO, Serrano JR, Jiang H, Roh J, Finn MB, Sullivan PM, Esparza TJ, et al: Age-Dependent Effects of apoE Reduction Using Antisense Oligonucleotides in a Model of beta-amyloidosis. Neuron 2017, 96:1013–1023 e1014.

74. Bennett ML, Bennett FC, Liddelow SA, Ajami B, Zamanian JL, Fernhoff NB, Mulinyawe SB, Bohlen CJ, Adil A, Tucker A, et al: New tools for studying microglia in the mouse and human CNS. Proc Natl Acad Sci U S A 2016, 113:E1738–1746.

