## Supplemental Figures for "*Ms4a4a* deficiency ameliorates plaque pathology in a mouse model of amyloid accumulation"

### Supplemental Figure 1

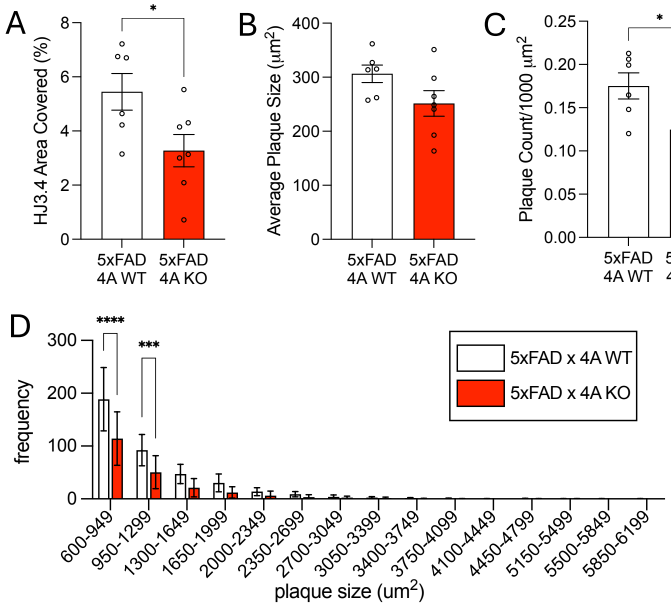

### Supplemental Figure 2

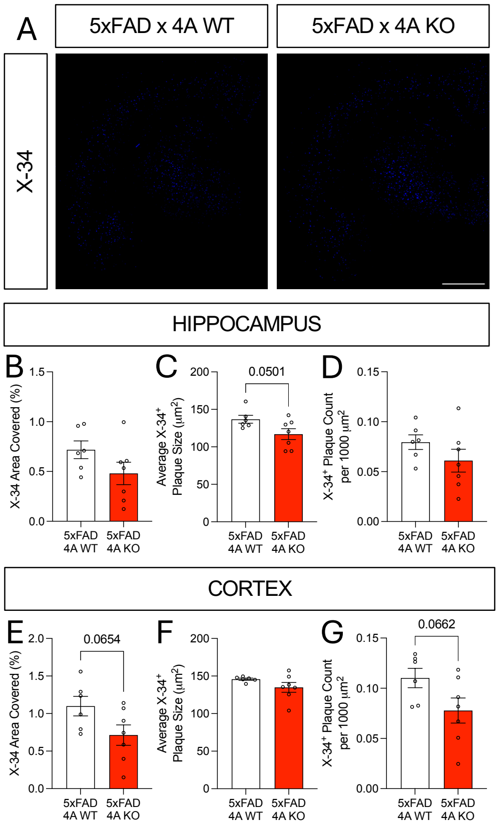

### Supplemental Figure 3

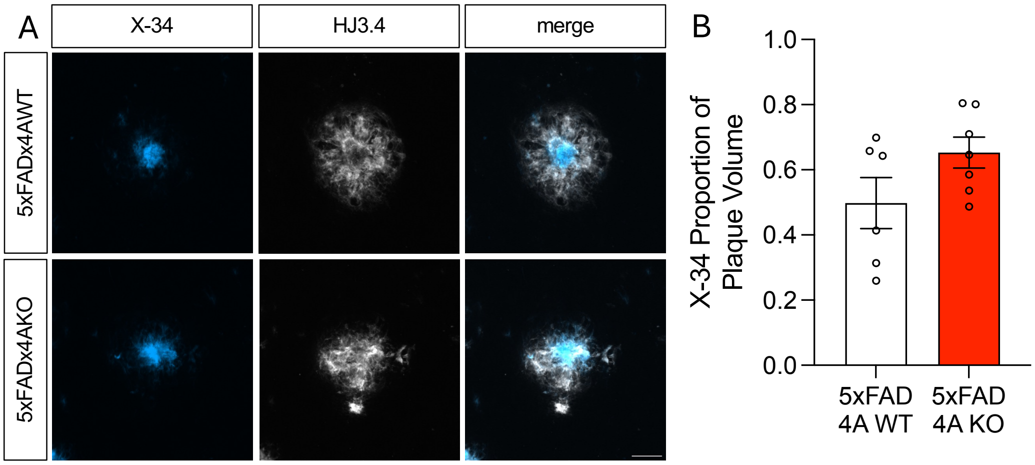

### Supplemental Figure 4

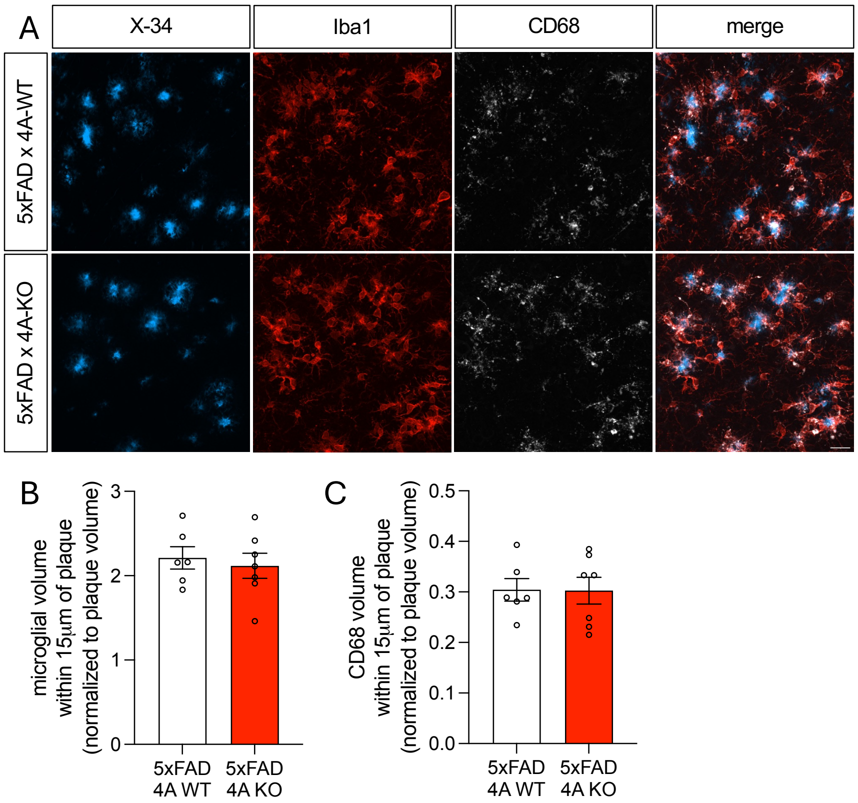

### Supplemental Figure 5

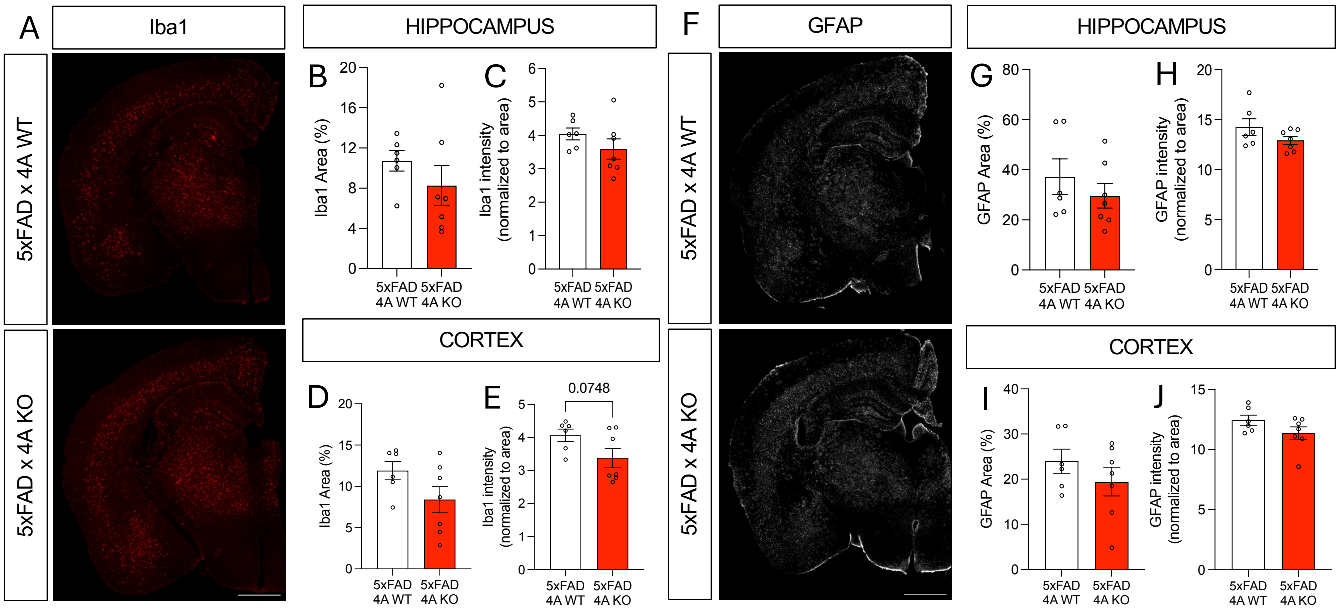
